# Choosing to view morbid information involves reward circuitry

**DOI:** 10.1101/795120

**Authors:** Suzanne Oosterwijk, Lukas Snoek, Jurriaan Tekoppele, Lara Engelbert, H. Steven Scholte

## Abstract

People often seek out stories, videos or images that detail death, violence or harm. Considering the ubiquity of this behavior, it is surprising that we know very little about the neural circuits involved in choosing negative information. Here we show that choosing intensely negative stimuli engages similar brain regions as those that support extrinsic incentives and “regular” curiosity. Participants made choices to view negative and positive images, based on negative (e.g., a soldier kicks a civilian against his head) and positive (e.g., children throw flower petals at a wedding) verbal cues. We hypothesized that the conflicting, but relatively informative act of choosing to view a negative image, resulted in stronger activation of reward circuitry as opposed to the relatively uncomplicated act of choosing to view a positive stimulus. Indeed, as preregistered, we found that choosing negative cues was associated with activation of the striatum, inferior frontal gyrus, anterior insula, and anterior cingulate cortex, both when contrasting against a passive viewing condition, and when contrasting against positive cues. These findings nuance models of decision-making, valuation and curiosity, and are an important starting point when considering the value of seeking out negative content.

---

Humans are active agents who often deliberately expose themselves to “morbid” information (e.g., information involving death, violence or harm). People choose to watch gruesome documentaries, click on links detailing terrifying attacks and visit locations of horrible events. Surprisingly, the fact that people experience curiosity for negative information, and often act on this feeling, is rarely addressed in theoretical models of curiosity and decision-making. Moreover, empirical work on this phenomenon is markedly limited and studies investigating the neural circuits involved in choosing negative information are virtually non-existent. Nevertheless, “morbid curiosity” is an important topic for investigation, because this ubiquitous behavior appears to be at odds with the idea that value and reward drive human information seeking. The present paper aims to expand the scientific enquiry of curiosity and choice, by investigating how the brain, and reward-related brain regions in particular, represent a deliberate choice to view intensely negative images that portray death, violence or harm.

In the last decades, much progress has been made in understanding the neuroscience of choice, valuation and curiosity. Yet, when studying choice, decision-making scientists typically focus on extrinsically rewarding stimuli, such as monetary rewards (Braver, Krug, Chiew, Kool, Westbrook, Clement et al., 2014). Decisions regarding intrinsically rewarding stimuli are targeted less frequently (Murayama, 2018) and even less is known about the neural representation of seeking negative information that, at first glance, does not seem to have reward value at all (Elliot, 2006). Similarly, in the field of curiosity - defined as an intrinsically motivated drive state for information (Kidd & Hayden, 2015; Golman & Loewenstein, 2015; Kobayashi, Ravaioli, Baranès, Woodford & Gottlieb, 2019) - research on curiosity for negative information is scarce (Murayama, 2018; see for exceptions Oosterwijk, 2017; Hsee & Ruan, 2016; Rimé, Delfosse & Corsini, 2005; Zuckerman & Litle, 1986). A handful of neuroscience studies have demonstrated that curiosity engages similar neural circuits as extrinsic reward, but only when examining positive or neutral material, such as trivia questions (e.g., Gruber, Gelman & Ranganath, 2014; Kang, Hsu, Krajbich, Loewenstein, McClure, Wang & Camerer, 2009). At present, it is thus unclear whether curiosity for “morbid” information is supported by similar neural mechanisms as those that support extrinsic incentives and “regular” curiosity.

Neuroscientific evidence suggests that curiosity, choice and reward are supported by a highly similar constellation of brain regions (Kidd & Hayden, 2015; Sakaki, Yagi & Murayama, 2018). Reward-related decision making, predominantly studied by focusing on monetary gains or losses, engages the dorsal striatum (caudate, putamen), ventral striatum (NAcc), orbitofrontal cortex (OFC), bilateral anterior insula, anterior cingulate cortex (ACC) dorsomedial prefrontal cortex/supplementary motor area (dmPFC/SMA) and frontal and parietal regions often associated with cognitive control (Levy & Glimcher, 2012; Liu, Hairston, Schrier & Fan, 2011; Samanez-Larkin & Knutson, 2015; Diekhof, Kaps, Falkai & Gruber, 2012; Bartra, McGuire & Kable, 2013). Several studies targeting curiosity demonstrated similar neural regions. For example, Kang and colleagues (2009) found increased activation in the inferior frontal gyrus (IFG), caudate and putamen when inducing curiosity by presenting trivia questions. When curiosity was relieved (i.e., when the answer to the question was given) they found engagement of the putamen and IFG. In another study targeting trivia questions, Gruber and colleagues (2014) found increased activation in the dorsal and ventral striatum and IFG for questions associated with high curiosity ratings. Other work has shown that the induction and relief of curiosity engages regions that are associated with salience detection and uncertainty (Singer, Critchley & Preuschoff, 2009; Menon & Uddin, 2010), including the anterior insula and ACC (Jepma, Verdonschot, van Steenbergen, Rombouts, & Nieuwenhuis, 2012; van Lieshout, Vandenbroucke, Muller, Cools & de Lange, 2018). In short, the limited work on curiosity so far, demonstrates that curiosity for relatively positive and neutral material engages neural regions that are also recruited during the computation of value and the anticipation of reward. Whether these regions also engage when people act on their curiosity for negatively valenced information is currently unknown.

In the present study, we tested the preregistered hypothesis that the striatum and IFG (ROI-analyses) and the anterior insula and ACC (whole-brain analyses) will engage more when people deliberately choose to view negative images, as compared to a passive viewing condition. This hypothesis reflects our assumption that “morbid curiosity”, expressed by a choice to view a negative stimulus, engages similar neural regions as regular curiosity (van Lieshout et al., 2018; Gruber et al., 2014; Jepma et al., 2012; Kang et al., 2009). In addition, we tested a second hypothesis that the regions described above will engage more strongly when people choose to view a negative stimulus, as compared to a positive stimulus. This hypothesis is based on our assumption that the informational value of negative images is relatively high. Compared to positive information, negative information may be more novel, rare, deviant, uncertain, challenging or complex (Unkelbach, Fiedler, Bayer, Stegmuller & Danner, 2008; Baumeister, Bratslavsky, Finkenauer, & Vohs, 2001) – these information characteristics engage reward circuitry and evoke curiosity (Kidd & Hayden, 2015; Sakaki et al., 2018; Berlyne, 1966; Kashdan & Silvia, 2009). Furthermore, a choice for negativity may involve a tradeoff between benefits (e.g., understanding something complex) and costs (e.g., being emotionally perturbed by a stimulus). Choosing a positive stimulus (e.g., viewing a family picnicking in the park), does not involve such costs, and may also have less benefits in terms of accessing novel, deviant or complex information. In this sense, choosing negativity (or “morbid curiosity”) is a conflict state; people want information, without predicting that they will like the information (see also Litman, 2005; Rimé et al., 2005). Previous work suggests that reward circuitry engagement is most pronounced when actions or decisions are ambiguous or unclear (Floresco, 2015). In line with this, we predict that the conflicting, but relatively informative act of choosing to view a negative image will paradoxically result in stronger activation of reward circuitry as opposed to the relatively uncomplicated act of choosing to view a positive stimulus.

We build upon previous work that demonstrates that people are interested in and fascinated by social negative images and prefer to view these stimuli above and beyond neutral alternatives (Oosterwijk, 2017; Oosterwijk, Lindquist, Adebayo & Barrett, 2016). We used an established choice paradigm (Oosterwijk, 2017) that presented participants with choices to view images that depicted social negative and positive situations, taken from validated affective picture databases. Importantly, this paradigm solely targets intrinsic motivation; participants were not financially rewarded for their choices. Moreover, this paradigm targets a behavioral expression of “wanting”, and not the extent to which people “like” the images (Litman, 2005; Berridge, Robinson & Aldridge, 2009). The active-choice condition was compared to a passive-viewing condition, using a yoked procedure that has been previously used to study responses to controllable and incontrollable stressors (Wood, Wheelock, Shumen, Bowen, Verhoef & Knight, 2016; Amat, Baratta, Paul, Bland, Watkins & Maier, 2005). In the present study, a yoked design allowed us to investigate the effect of choice, while controlling for general affective, semantic and visual processing.

In the choice condition, people were presented with verbal cues, describing negative images (e.g., rescue workers treat a wounded man; a soldier kicks a civilian against his head) and positive images (e.g., children throw flower petals at a wedding; partying people carry a crowd surfer). The presentation of the verbal cue was labeled the induction phase (see Figure 1). Following the cue, participants chose whether they wanted to see the image corresponding to the description, or not. In the relief phase (see Figure 1), participants viewed the corresponding image when responding yes and a blurred version when responding no. The passive-viewing condition was fully yoked to the active-choice condition. In other words, each participant in the passive-viewing condition did not make choices, but was confronted with the choice profile of a participant in the active-choice condition. Participants were first exposed to the cue, then they confirmed the choice that was determined by the computer (to mirror the motor response made in the choice condition), then they viewed either the full image or a blurred version. Importantly, this yoked design isolates the psychological process that we aim to investigate (i.e., a deliberate choice to view a stimulus), while keeping all other factors constant (i.e., cues and images). In line with curiosity theories that argue that exploratory behavior is an important component of curiosity (Litman, 2005; Kashdan & Silvia, 2009; Loewenstein, 1994), we propose that the potential to choose will make participants’ subjective experience of curiosity more salient. In other words, participants’ subjective state of curiosity may be more at the fore-front of consciousness in the active-choice condition (because it will inform participants’ decisions) than in the passive-viewing condition.

**Figure 1.**
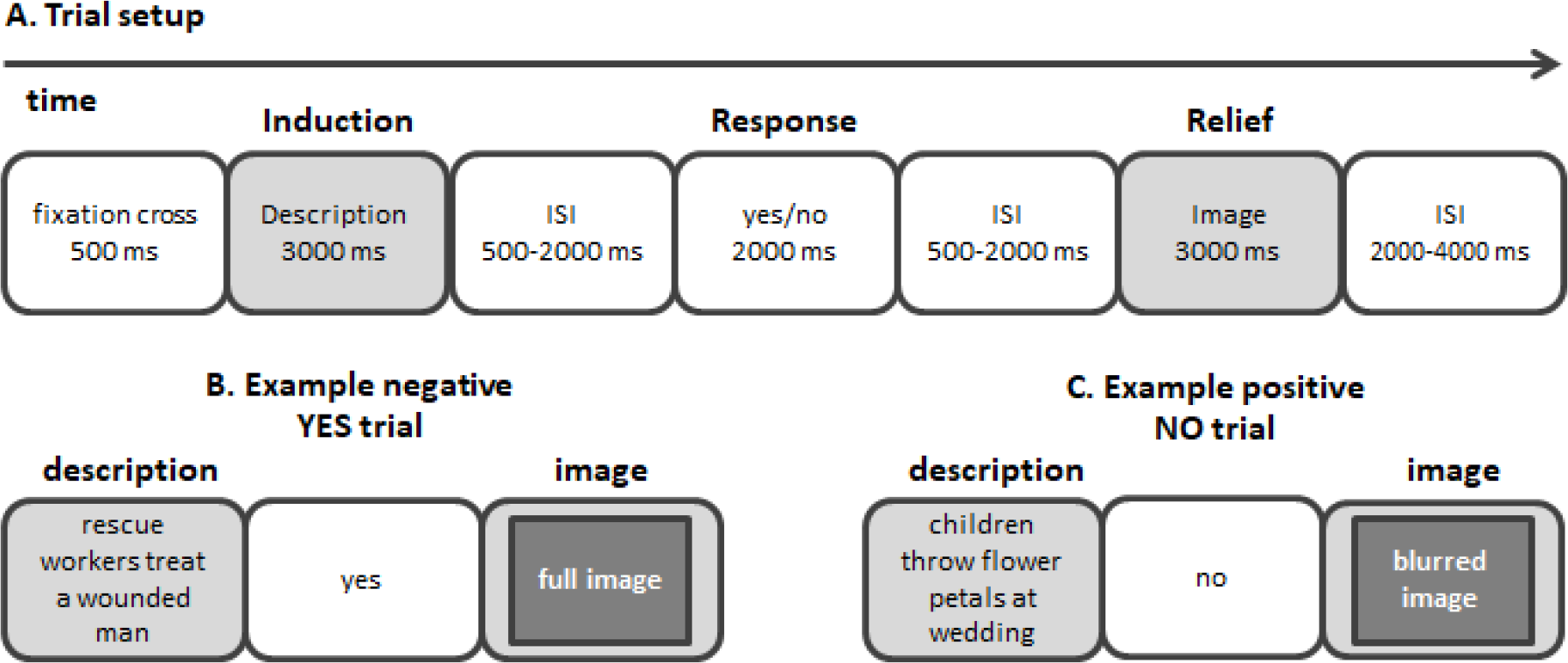
Overview of paradigm. **A)** The setup of the trials in the choice-condition and passive-viewing condition. Note that in the active-choice condition, participants chose whether they wanted to see the image corresponding to the description during the yes/no response event. In the passive-viewing condition participants did not choose, but confirmed the decision seemingly determined by the computer during the yes/no response event. **B)** An example of a negative description and the consequence of a yes response (either given by the participant, or determined by the computer). **C)** An example of a positive description and the consequence of a no response (either given by the participant, or determined by the computer).

We selected positive and negative images from the International Affective Picture System database (IAPS; Lang, Bradley & Cuthbert, 2008) and the Nencki Affective Picture System database (NAPS; Marchewka, Żurawski, Jednoróg & Grabowska, 2014; see method section for more details). Importantly, we matched negative and positive images in terms of valence extremity to ensure that, on average, positive images were perceived as equally positive as negative stimuli were perceived negative. A similar procedure was performed for the positive and negative descriptions (i.e., cues; see method section for further details).

Our scanning protocol was performed on a 3T scanner. For all details on image acquisition, preprocessing and first- and second-level analyses, please see the method section. Our analysis protocol held confirmatory and exploratory analyses. Hypotheses and corresponding contrasts for the confirmatory analyses, exclusion criteria, ROIs and corrections for multiple comparisons were preregistered on the Open Science Framework, prior to data analysis (osf.io/gdtk9, Oosterwijk, 2017).

The first-level model included eight predictors to capture the research design: 2 (phase: induction phase vs. relief phase) × 2 (choice: yes, full image vs. no, blurred image) × 2 (valence: negative vs. positive). Additionally, we added a single predictor for the motor response associated with the decision/confirmation response and six motion predictors based on estimated motion correction parameters. First level contrasts only involved trials associated with yes choices (i.e., full image trials); trials associated with no responses (i.e., blurred image trials) were not used for further group level analysis.

In the ROI-based analyses, we focused on voxels within two a-priori defined ROIs: bilateral striatum and bilateral inferior frontal gyrus (IFG). The ROIs were based on the Harvard-Oxford Subcortical Atlas (striatum; caudate, putamen and nucleus accumbens) and the Harvard-Oxford Cortical Atlas (IFG; pars opercularis and pars triangularis) with a threshold for probabilistic ROIs > 0 (Craddock, James, Holtzheimer & Mayberg, 2012). In the group-level analyses targeting these two ROIs, we calculated two contrasts that reflected our confirmatory hypotheses, separately for the induction and relief phase. We hypothesized stronger activation in the ROIs when participants processed a negative cue/image in the active-choice condition as compared to that same event in the passive-viewing condition (i.e., (*β*_neg | active_ − *β*_neg | passive_) > 0). In addition, we hypothesized stronger activation in the ROIs when participants processed a negative cue/image in the active-choice condition as compared to a positive cue/image in the active-choice condition, controlling for passive viewing (i.e., (*β*_neg | active_ − *β*_neg | passive_) − (*β*_pos | active_ − *β*_pos | passive_) > 0). For these confirmatory ROI analyses, we used nonparametric permutation-based inference in combination with Threshold-Free Cluster Enhancement (TFCE; Smith & Nichols, 2009) as implemented in *FSL randomise* (Winkler, Ridgway, Webster, Smith, & Nichols, 2014) and thresholded voxelwise results at *p* < 0.025 (correction for two ROIs). Note that this analysis allows for voxel-wise inference (i.e., no cluster-based correction is used). In addition to the confirmatory ROI analysis, we conducted an exploratory whole-brain group-level analysis. In addition to the two confirmatory contrasts mentioned in the previous section, we tested three exploratory contrasts, separately for the induction and relief phase with a voxel-wise *p*-value threshold of 0.005 and a cluster-wise *p*-value of 0.05). Full details regarding the exploratory analyses can be found in the method section.

## Results

### Participants

In total, sixty participants signed informed consent and underwent our scanning protocol. We implemented a preregistered eligibility criterion that only participants who chose negative and/or positive images in 40% or more of the trials would be paired with a participant in the passive-viewing condition. Neuroimaging analyses were performed on a sample of 54 participants (38 women; *M*_age_ = 22.4, *SD* = 2.9); with 27 participants in the active-choice condition and 27 participants in the passive-viewing condition.

### Behavior and subjective report

In the active-choice condition, participants chose to view the negative image in 80.6% of the trials; participants chose to view the positive image in 94.8% of the trials. In the active-choice condition, participants reported that they followed their curiosity more when making choices for negative cues (*M* = 6.20, *SD* = .71) as compared to positive cues (*M* = 4.72, *SD* = 1.82), *t*(24) = 3.49, *p* = .002, 95% CI [0.60, 2.36], *d_z_* = .70. Similarly, participants expressed more curiosity for negative cues (*M* = 5.41, *SD* = 1.28) than for positive cues (*M* = 4.44, *SD* = 1.60), *t*(27) = 2.42, *p* = .023, 95% CI [0.15, 1.78], *d_z_* = .47, in the passive-viewing condition. This finding is consistent with previous results that people find negative social information generally more interesting and fascinating than positive social information (Oosterwijk, 2017; Oosterwijk et al., 2016). As expected, endorsed ratings on the curiosity question were significantly higher in the active-choice condition as compared to the passive viewing condition, but only for negative cues (*M* = 6.20 vs. *M* = 5.41), *t*(50) = 2.74, *p* = .009, 95% CI [0.22, 1.36], *d* = .77, and not for positive cues (*M* = 4.72 vs. *M* = 4.44), *t*(50) = .58, *p* = .563, 95% CI [−0.68, 1.23], *d* = .16.

### ROI analyses

Our first set of hypotheses focused on contrasting neural activity when participants processed a negative cue in the active-choice condition with that same event in the passive-viewing condition (i.e., negative _active > passive_). As predicted, a confirmatory ROI analysis demonstrated more activation in the striatum when participants viewed a negative cue that was chosen (in the active-choice condition) as compared to watching that same negative cue in the passive-viewing condition. Figure 2 shows that this contrast produced significant voxels across the striatum, in the caudate, putamen and NAcc. The ROI analysis targeting the IFG also demonstrated stronger activation in the active-choice condition as compared to the passive-viewing condition, further confirming our hypotheses.

**Figure 2.**
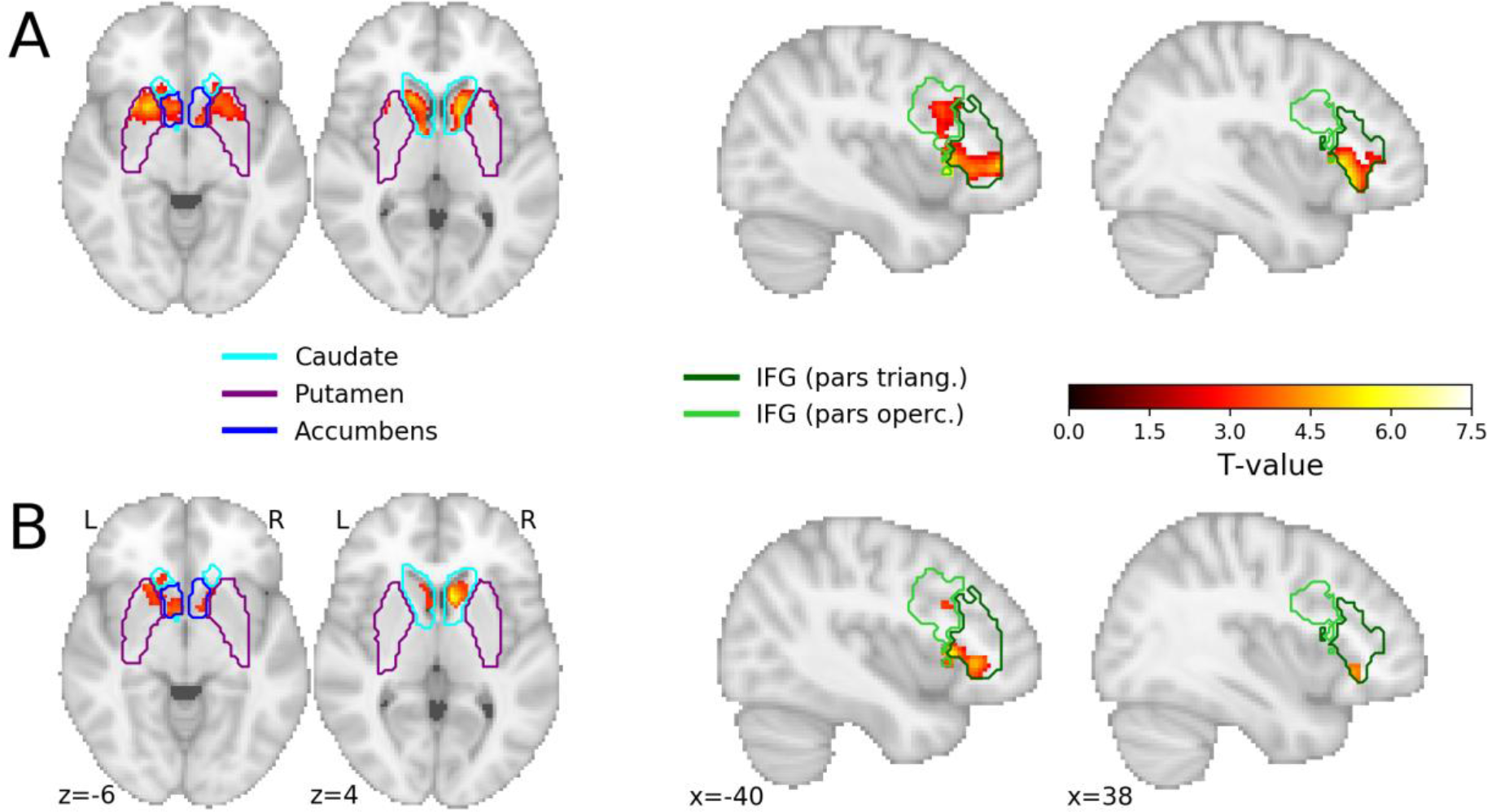
Results of confirmatory ROI analyses for the induction phase **A**) the contrast negative _active > passive_ **B**) the contrast negative _active > passive_ > positive _active > passive_. Voxels in red/yellow represent significant *t*-values (*p* < 0.05, corrected for multiple comparisons using the maximum statistic approach). The colored outlines represent the different brain regions within the probabilistic ROIs for the striatum (left) and inferior frontal gyrus (IFG; right). The outlines represent the border of the ROIs thresholded at 0. When voxels within one ROI had a nonzero probability in more than one brain region (e.g., the caudate and nucleus accumbens), the voxel was assigned to the brain region with the largest probability.

Our second set of hypotheses focused on comparing a choice for negative information with a choice for positive information, controlling for general semantic, affective and visual processing (i.e., negative _active > passive_ > positive _active > passive_). As predicted, a confirmatory ROI analysis demonstrated more activation in the striatum when participants viewed a negative cue that was chosen as compared to a positive cue that was chosen, relative to watching that same negative or positive cue in the passive-viewing condition. Again, significant voxels were found across the striatum, in the caudate, putamen and NAcc. A similar effect was found in the IFG (See Figure 2). To explore the directionality of the effects, we extracted the beta-weights for the individual regressors for both ROIs. A visual inspection of plotted weights suggest that the patterns of neural activation reported for the striatum, are driven both by activation in the striatum when viewing negative cues in the active-choice condition, and deactivation in the striatum when viewing negative cues in the passive-viewing condition. Further details can be found in Figure S1 of the Supplementary Materials.

### Whole-brain analyses

In addition to the confirmatory analyses reported above, we performed a whole-brain analysis (cluster-corrected with a voxel-wise threshold of *p* < .005 and a cluster-wise threshold of *p* < .05) for the two confirmatory contrasts (see Figure 2). In addition to activation in the regions targeted in the confirmatory ROI analyses, the whole-brain analyses for the negative _active > passive_ contrast demonstrated robust activation in the ACC, paracingulate gyrus, superior frontal gyrus, middle frontal gyrus, OFC, insular cortex, frontal operculum, frontal pole, temporal pole, thalamus and brain stem, when participants viewed a negative cue that was chosen (in the active-choice condition) as compared to watching that same negative cue in the passive-viewing condition. A complete table of the significant clusters can be found in the Supplementary Materials (Table S2, as well as the significant clusters associated with the positive _active > passive_ contrast, Table S3). The whole-brain results for the negative _active > passive_ > positive _active > passive_ contrast are presented in Table 1. This contrast demonstrated stronger activation in the ACC, paracingulate gyrus, superior frontal gyrus, OFC, insular cortex and frontal operculum, when participants viewed a negative cue that was chosen as compared to a positive cue that was chosen (relative to watching that same negative or positive cue in the passive-viewing condition).

**Table 1.** Cluster statistics and associated brain regions from the exploratory whole-brain analysis of the negative _active > passive_ > positive _active > passive_ contrast. The *X*, *Y*, and *Z* coordinates refer to MNI152 (2mm) space. The regions are taken from the Harvard-Oxford (sub)cortical atlas (Craddock et al., 2012) and voxels are assigned to regions based on their maximum probability across all ROIs within the atlas. *K* refers to the number of voxels within a particular region.

| Cluster nr. | Cluster size | Cluster max. | X | Y | Z | Region | K | Max. |
| --- | --- | --- | --- | --- | --- | --- | --- | --- |
| 1 | 3158 | 4.82 | -8 | 30 | 24 | Left Paracingulate Gyrus | 831 | 4.58 |
|  |  |  |  |  |  | Right Paracingulate Gyrus | 741 | 4.70 |
|  |  |  |  |  |  | Left Superior Frontal Gyrus | 657 | 4.64 |
|  |  |  |  |  |  | Right Cingulate Gyrus, anterior division | 317 | 4.42 |
|  |  |  |  |  |  | Left Cingulate Gyrus, anterior division | 300 | 4.82 |
|  |  |  |  |  |  | Right Superior Frontal Gyrus | 201 | 3.94 |
|  |  |  |  |  |  | Left Juxtapositional Lobule Cortex | 76 | 3.27 |
| 2 | 1928 | 4.84 | -42 | 20 | 4 | Left Frontal Orbital Cortex | 602 | 4.64 |
|  |  |  |  |  |  | Left Insular Cortex | 234 | 4.17 |
|  |  |  |  |  |  | Left Inferior Frontal Gyrus, pars triangularis | 229 | 4.17 |
|  |  |  |  |  |  | Left Frontal Operculum Cortex | 205 | 4.84 |
|  |  |  |  |  |  | Left Inferior Frontal Gyrus, pars opercularis | 93 | 3.65 |
|  |  |  |  |  |  | Left Temporal Pole | 57 | 4.31 |
|  |  |  |  |  |  | Left Subcallosal Cortex | 29 | 3.22 |
|  |  |  |  |  |  | Left Caudate | 121 | 3.48 |
|  |  |  |  |  |  | Left Putamen | 110 | 3.86 |
|  |  |  |  |  |  | Left Thalamus | 48 | 3.84 |
|  |  |  |  |  |  | Left Accumbens | 39 | 3.52 |
| 3 | 692 | 4.64 | 36 | 24 | -16 | Right Frontal Orbital Cortex | 443 | 4.64 |
|  |  |  |  |  |  | Right Insular Cortex | 79 | 3.80 |
|  |  |  |  |  |  | Right Frontal Operculum Cortex | 70 | 3.39 |
|  |  |  |  |  |  | Right Temporal Pole | 44 | 3.40 |
|  |  |  |  |  |  | Right Inferior Frontal Gyrus, pars triangularis | 30 | 3.27 |
| 4 | 464 | 4.74 | 10 | 14 | 4 | Right Caudate | 265 | 4.74 |
|  |  |  |  |  |  | Right Putamen | 26 | 3.49 |
| 5 | 296 | 3.81 | -36 | 12 | 26 | Left Middle Frontal Gyrus | 111 | 3.72 |
|  |  |  |  |  |  | Left Inferior Frontal Gyrus, pars opercularis | 111 | 3.81 |
|  |  |  |  |  |  | Left Precentral Gyrus | 64 | 3.46 |

To interpret the whole brain results further, we used the “decoder” function from Neurosynth.org (Yarkoni, Poldrack, Nichols, Van Essen, & Wager, 2011) to find key terms associated with particular patterns of neural activation. The neural pattern produced by the negative _active > passive_ > positive _active > passive_ contrast resulted in the following key terms (top-10): reward, task, monetary, semantic, anticipation, incentive, demands, fear, autobiographical, retrieval (see further Supplementary Materials, Table S1). Although it is important to be careful with drawing reverse inference conclusions about the psychological meaning of neural activation (Poldrack, 2006), these terms at minimum suggest that the neural pattern associated with choosing negative content (relative to choosing positive content and passive-viewing) is similar to neural patterns associated with reward and the processing of extrinsic incentives.

It is important to note that none of the confirmatory ROI analyses nor any of our exploratory analyses showed significant differences in the relief phase (i.e., when viewing the image). This is in line with previous work on curiosity (Gruber, et al., 2014) that found robust neural activation when inducing curiosity (e.g., presentation of trivia questions), but not when relieving curiosity (e.g., presentation of trivia answers). Other work on curiosity, that did find differences in the relief phase, contrasted a condition in which curiosity was relieved, with a condition that withheld information (van Lieshout et al., 2018) or a condition that showed irrelevant information (Jepma et al., 2012). In the present study, however, we contrasted the relief phase in the active-choice condition with viewing the exact same information in the passive-viewing condition. Since we found that participants in the passive-viewing condition also reported a reasonable amount of curiosity in response to the cue, curiosity relief may have occurred in both the active-choice and the passive-viewing condition.

**Figure 3.**
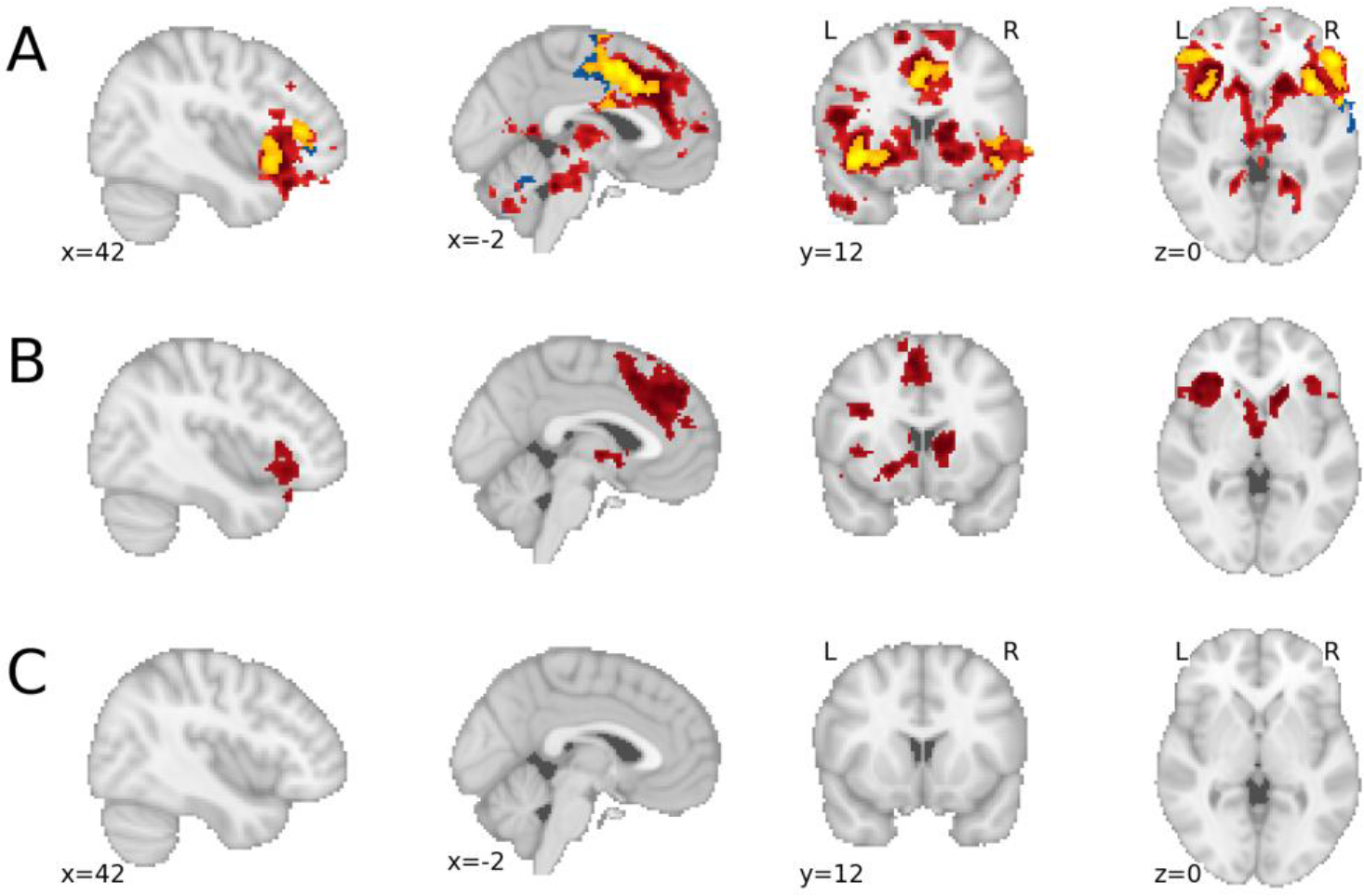
Results of the exploratory whole-brain analyses for the induction phase: **A**) the contrast negative _active > passive_ (red), positive _active > passive_ (blue), and their conjunction (yellow); **B**) the contrast negative _active > passive_ > positive _active > passive_; **C**) the contrast positive _active > passive_ > negative _active > passive_ (empty).

## Discussion

In the last decade, neuroscientific research has made a major contribution to a better understanding of choice, value and curiosity. Most of this work, however, has focused on extrinsically rewarding information (Bartra et al., 2013; Braver et al., 2014), or on an intrinsically-motivated curiosity for neutral or positive information (e.g., Kang et al., 2009; van Lieshout et al., 2018; Gruber et al., 2014). The present study demonstrates that choosing intensely *negative* stimuli engages similar brain regions as those that support extrinsic incentives and regular curiosity (Diekhof et al., 2012; Bartra et al., 2013; Kidd & Hayden, 2014; Sakaki et al., 2018). We found that a deliberate choice for death, violence or harm is associated with activation in the striatum (NAcc, caudate and putamen), inferior frontal gyrus, anterior insula, orbitofrontal cortex and anterior cingulate cortex. This pattern was present both when we contrasted negative cues (that were chosen) with passively viewing these cues, and when we contrasted negative cues (that were chosen) with positive cues (that were chosen), controlling for passive viewing. These findings reflect the induction phase in which participants anticipated on their choice; we found no differences in the relief phase in which participants viewed images.

Although we found activation in regions associated with reward and incentives (Diekhof et al., 2012; Bartra et al., 2013), it is important to be careful with the conclusion that it is “rewarding” to choose negative information (Poldrack, 2006). Reward regions are associated with a multitude of psychological processes (Bartra et al., 2013), and reward is a construct with different dissociable dimensions, including “liking” and “wanting” (e.g., Weber, Kahnt, Quednow & Tobler, 2018; Berridge, Robinson & Aldridge, 2009). In addition, the distributed pattern of activation that we found most likely reflects psychological processes beyond reward as well. In the next section, we address several, non-mutually exclusive, interpretations of this distributed pattern, including an interpretation regarding the potential reward value of negativity, the possibility that this neural pattern reflects uncertainty, and an interpretation that considers the particular characteristics of the task.

Golman and Loewenstein (2015) propose that a desire to obtain information can be driven by a motive to make better decisions, a motive to experience pleasantness, or a motive to engage with information “*for its own sake*” (Golman & Loewenstein, 2015, pp. 3). The negative choices made by our participants are most likely consistent with the latter intrinsic motive. Participants chose images that were not hedonically pleasing and, in contrast to other recent (neuroscientific) studies targeting curiosity (e.g., Kobayashi et al., 2019; van Lieshout et al., 2018), there was no monetary outcome associated with choosing (negative) images. The question that follows then is: what is the value of negativity?

One possibility is that knowledge acquisition is inherently valuable (Kidd & Hayden, 2014; Murayama, FitzGibbon & Sakaki, 2019; Marvin & Shohamy, 2016), even when people acquire knowledge about negative social situations that involve death, violence or harm. Indeed, the knowledge update that follows from engaging with negative information might be valuable for building a realistic model of the world (see Baumeister, et al., 2001) or for dealing with future aversive situations. Applying this interpretation to our findings, we propose that the brain predicts a larger information update in the negative choice condition than in the positive choice condition, given that negative information is relatively complex, challenging and deviant from the norm (Unkelbach et al., 2008). This is reflected in stronger activation of reward circuitry (e.g., NAcc, caudate, putamen) that might track the expected value of knowledge acquisition, or the salience of the information (Bartra et al., 2013). Note that we focus here on the informational value of valenced stimuli in epistemic terms (Berlyne, 1966; Loewenstein, 1994; Litman, 2005; Murayama et al., 2019; see also Tamir, 2016), but the value of choosing to engage with negative or positive stimuli may also lie in the emotional experiences or sensations that are evoked by the stimulus (Zuckerman, 1979; Zuckerman & Litle, 1986).

Another possible explanation for the present findings lies in the momentary uncertainty that people may experience when viewing negative cues (e.g., How extreme will the image be?), in combination with the predicted reduction of this uncertainty when choosing to view images. Curiosity is often seen as a desire to resolve uncertainty (Berlyne, 1966; Loewenstein, 1994; Gottlieb & Oudeyer, 2018) and several recent studies have demonstrated that people experience higher levels of curiosity for uncertain stimuli (Kobayashi et al., 2019; van Lieshout, et al., 2018). People even engage with aversive stimuli to reduce uncertainty, preferring a reduction in uncertainty above a negative outcome (e.g., shock, Hsee & Ruan, 2016). Neuroscience studies have shown that uncertainty is associated with activation in the OFC, ACC and anterior insula (e.g., Bach & Dolan, 2012; Harris, Sheth & Cohen, 2008; Singer et al., 2009), regions we also found to be active in the active-choice condition. Moreover, we found that the OFC, ACC and anterior insula were more strongly engaged when viewing chosen negative cues, as compared to chosen positive cues. The neural pattern found in the negative choice condition (as compared to the positive choice condition) may thus reflect a higher level of outcome uncertainty, and/or a stronger expected reduction in uncertainty, in interaction with, or irrespective of (Gottlieb & Oudeyer, 2018), the reward value of the information.

A final interpretation of the present findings, in particular regarding the engagement of the ACC, revolves around the demands of the task that people performed in the choice-condition. Recently, it has been suggested that the dACC may be particularly active in effortful, complex or exploratory tasks that demand cognitive control, as compared to tasks that can be performed by engaging in automatic behavior (Shenhav, Cohen & Botvinick, 2016). When applying this perspective to the current findings, stronger ACC activation for negative as opposed to positive choice may be explained by the relatively complex cost-benefit analysis that precedes a choice to choose a negative stimulus, as compared to the relatively automatic or “default” decision to choose a positive stimulus. This interpretation is consistent with the characterization of “morbid curiosity” as a conflict state, in which people “want” information that they do not “like” (see also Litman, 2005; Rimé et al., 2005).

The present study has a few limitations that we should address. First of all, all contrasts are relative to the passive-viewing condition (as preregistered). We deliberately made this decision, because we wanted to isolate the neural activation associated with a deliberate choice to view a stimulus. More specifically, in a direct contrast between chosen negative cues and chosen positive cues, it would have been impossible to know whether the pattern of neural activation was driven by *choosing* negative versus positive information, or by simply *viewing* negative versus positive cues. The yoked procedure, and the resulting contrasts, control for the latter, since activation associated with viewing negative vs. positive cues is subtracted out. Furthermore, confronting participants with emotional material that they cannot control, is common practice when scientists study affective/emotional experience (Lindquist, Satpute, Wager, Weber, & Barrett, 2016; Lang & Bradley, 2010) and emotion regulation (Buhle, Silvers, Wager, Lopez, Onyemekwu, Kober et al., 2014; Wager, Davidson, Hughes, Lindquist & Ochsner, 2008), and thus serves as a meaningful control condition. Nevertheless, it is important to note that our results in the striatum were partly driven by deactivation in the passive viewing condition (see Figure S1, Supplementary Materials). Since this pattern of deactivation was not predicted in our preregistered analysis protocol, and our study was not designed for optimal detection of directional effects, we will not discuss this finding further. A design that contrasts choosing versus passive viewing within-subjects may clarify whether explicitly anticipating a negative outcome that cannot be controlled indeed deactivates the striatum.

A second limitation is that the present study focused on behavior, without incorporating trial-by-trial ratings of curiosity. This restricts the extent to which the present findings speak to the subjective experience of curiosity. In regular curiosity, for example when processing trivia questions, there are little costs associated with acting on curiosity, and thus it may be sufficient to focus on subjective ratings of curiosity. With morbid curiosity, however, the stakes are higher. Although people can experience curiosity for a negative stimulus without choosing to engage with it, behavior is, in our opinion, the most straightforward indicator of how the conflict state of morbid curiosity is resolved (Litman, 2005; Rimé et al., 2005). In other words, only when people choose to engage with negative information can we deduce that the predicted benefits (e.g., knowledge acquisition, uncertainty reduction) outweigh the predicted costs of engaging with the information (e.g., not being able to cope with the content). Furthermore, a focus on choice connects to the many behavioral expressions of this phenomenon in the real world (e.g., “rubbernecking” on the freeway; clicking on a social media link). Future research should investigate whether the subjective experience of curiosity for negative stimuli is associated with a similar neural pattern (e.g., reward circuitry, ACC, insula, OFC) as the pattern found with the present choice paradigm.

Despite the questions that the present study provokes, our findings represent an important step in nuancing models of decision-making, valuation and curiosity. In light of the ubiquity of exploring negativity in daily life, we believe that it is crucial to start thinking about the value of seeking out negative content.

## Methods

### Participants

Participants consisted of a convenience sample of students at the University of Amsterdam. The budget allowed for scanning of a maximum of 60 participants. After applying the exclusion criteria, the total sample consisted of 54 participants, including 38 women (*M*_*age*_ = 22.4, *SD* = 2.9) and 16 men (*M*_*age*_ = 23.8, *SD* = 1.8).

### Design

This study used a 2 (choice: active-choice vs. passive-viewing) × 2 (phase: induction vs. relief) × 2 (valence: negative vs. positive) mixed design. The variable choice was varied between participants and consisted of an active-choice condition and a passive-viewing condition. The variable phase was varied within participants and reflected the presentation of the cue (i.e., induction) vs. the presentation of the image (i.e., relief). The variable valence was varied within participants and reflected the negative vs. positive content of the cues/images.

### Materials

#### Experimental task

The present study utilized a choice task (Oosterwijk, 2017) that presented participants with verbal cues describing negative and positive images, and offered them a choice to see these images or blurred versions. We used a yoked design that isolated the effects of choice, controlling for general affective, semantic and visual processing (see also Wood, et al., 2016; Amat et al., 2005). This yoked design resulted in two conditions: the active-choice condition and the passive-viewing condition. Participant in the passive-viewing condition did not make choices, but were confronted with the choice profile of a yoked participant in the choice condition. Tasks in both conditions were programmed in Neurobs Presentation (https://www.neurobs.com/presentation). Behavioral data preprocessing was done using Python 3.5 and analyzed using IBM SPSS Statistics 22.0.

In the active-choice condition, participants were presented with 35 negative cues (e.g., rescue workers treat a wounded man; a soldier kicks a civilian against his head) and 35 positive cues (e.g., children throw flower petals at a wedding; partying people carry a crowd surfer) that described images, in random order. In each trial participants could choose, based on the cue, whether they wanted to view the corresponding image or not. The choice task consisted of a total of 70 trials. Each trial started with a fixation cross, presented for 500 ms, followed by the cue, presented for 3000 ms. The presentation of the cue was labelled as the induction phase (see also, Jepma et al., 2012; van Lieshout et al., 2018). The cue was followed by a jittered interval varying between 500-2000 ms. Subsequently, participants saw the words ‘yes’ and ‘no’ on the screen, and chose whether they wanted to see the image that was described by the cue, or not, by pressing one of two pre-specified buttons. Immediately following their response, the word ‘yes’ or ‘no’ turned green, indicating that their response was registered. Participants had a 2000 ms. time window to make their choice. If they had not made a choice after 2000 ms, the choice was automatically set to ‘no’. The response phase was followed by a jittered interval varying between 500-2000 ms. The interval was followed by the relief phase, in which the participants were presented with the image (size) when they chose ‘yes’. When participants chose not to see the corresponding image, they were presented with a blurred version of the image that was unrecognizably distorted (filter). Both the image and the blurred image were presented for 3000 ms. The relief phase was followed by a jittered inter-trial-interval varying between 2000-4000 ms. For a visual representation of the paradigm, please see Figure 1.

In the passive-viewing condition, participants were presented with the choice profile of a participant in the active-choice condition (i.e., the exact pattern of ‘yes’ and ‘no’ responses to the positive and negative cues for each participant in the active-choice condition was saved, and then re-used once as the computer generated pre-determined choice pattern for a participant in the passive-viewing condition). Participants were told in the introduction to the study that the computer would determine which images would be shown. The trial setup was identical to the active-choice condition, except for the following aspect. After participants were presented with the cue, the word ‘yes’ or ‘no’ turned green, indicating the choice of the computer. Participants were asked to confirm the choice made by the computer by pressing one of two pre-specified buttons, to mirror the motor response made in the active-choice condition.

The active-choice condition came with a filling problem: because participants could choose whether they wanted to view an image or not, some participants would see many more images than others. This filling problem can pose problems for modelling the BOLD response, due to lower efficiency in estimating contrasts for one subject over the other. Based on individual differences in choosing to view social negative information (Oosterwijk, 2017), we formulated an a-priori defined and preregistered eligibility criterion that only participants in the active-choice condition who chose negative and/or positive images in 40% or more of the trials (14/35 stimuli) would be paired with a subject in the passive-viewing condition. Based on this criterion five out of 33 participants in the active-choice condition were excluded from the sample. One other participant was excluded, because the functional scan was stopped prematurely. This resulted in 27 participants in the active-choice condition. The choice profiles of these 27 participants were yoked with 27 participants in the passive-viewing condition.

#### Cues

Cues were written to describe positive and negative images in one sentence. In a pilot study the cues were rated on valence (0 = negative to 100 = positive), and arousal (0 = low arousal to 100 = high arousal). Negative cues were rated more negatively than positive cues (*M* = 20.69, *SD* = 8.42 vs. *M* = 77.99, *SD* = 4.49), *t*(68) = −35.51, *p* < .001, and more arousing than positive cues (*M* = 68.45, *SD* = 6.38 vs. *M* = 28.93, *SD* = 5.99), *t*(68) = 26.71, *p* < .001. To check whether negative and positive cues were matched in terms of valence extremity we calculated mean-centered valence scores. An analysis of these scores demonstrated that, on average, positive cues were perceived as equally positive (*M* = 29.31, *SD* = 8.43) as negative stimuli were perceived as negative (*M* = 27.99, *SD* = 4.49), *t*(68) = .82, *p* = .417.

#### Images

Images were selected from the International Affective Picture System (IAPS; Lang, Bradley, Cuthbert, 2008) and the Nencki Affective Picture System (NAPS; Marchewka, Żurawski, Jednoróg & Grabowska, 2014); image codes are presented in the Supplementary Materials (Table S4). We selected negative images that portrayed situations of interpersonal violence, or social scenes involving a dead body or a harmed person. Negative images were selected when they had a valence rating below 4 (on a scale from 1 = negative to 9 = positive) and an arousal rating above 4.5 (on a scale from 1 = not arousing to 9 = extremely arousing). We selected positive images that portrayed joyful, loving or exciting interpersonal interactions. Positive images were selected when they had a valence rating above 6 (on a scale from 1 = negative to 9 = positive) and an arousal rating above 3 (on a scale from 1 = not arousing to 9 = extremely arousing). Negative and positive images differed significantly in terms of valence (*M* = 2.58, *SD* = .53 vs. *M* = 7.43, *SD* = .36), *t*(68) = −44.88, *p* < .001, and arousal (*M* = 6.18, *SD* = .75 vs. *M* = 4.78, *SD* = .90), *t*(68) = 7.07, *p* < .001. To check whether negative and positive images were matched in terms of valence extremity we calculated mean-centered valence scores. An analysis of these scores demonstrated that, on average, positive images were perceived as equally positive (*M* = 2.42, *SD* = .53) as negative stimuli were perceived as negative (*M* = 2.44, *SD* = .37), *t*(68) = −.22, *p* = .824.

#### Questionnaires

After the scanning session was completed, participants filled in the ‘Morbid curiosity in daily-life’ questionnaire (Oosterwijk, 2017) and the Dutch version of the Interpersonal Reactivity Index (IRI; De Corte, Buysse, Verhofstadt, Roeyers, Ponnet, & Davis, 2007). A short exit questionnaire asked participants two questions regarding the task they performed in the scanner. Participants in the active-choice condition were asked to rate to what extent they followed their curiosity when making choices for negative cues, and when making choices for positive cues, on a 1 (not at all) to 7 (very much) point scale. Participants in the passive-viewing condition were asked to rate to what extent they were curious about the negative cues, and the positive cues, on a 1 (not at all) to 7 (very much) point scale. The exit questionnaire concluded with demographic questions.

### Procedure

After signing the informed consent form, each participant received a thorough instruction. The active-choice condition was introduced as a study on how the brain represents choice. Participants were explained how they could make their choice, that they would always see the image of their choice, and that there were no right or wrong answers. Furthermore, participants were presented with an example of a negative and a positive cue, combined with the corresponding full image and blurred image, so that they knew what to expect when choosing the yes or no option. No mention was made of curiosity in the instruction. The passive-viewing condition was introduced as a study on the brain processes involved in reading image descriptions and viewing images. Participants were explained that the computer determined whether a description would be followed by a corresponding image. As in the active-choice condition, participants were presented with an example of a negative and a positive cue, combined with the corresponding full image and blurred image, so that they knew what to expect when the computer determined the yes or no option.

When comfortable and instructed, a structural T1-weighted anatomical scan was made. Then the participant performed the choice task or the passive task during fMRI acquisition in the scanner. After the scanning session, the participant filled in the questionnaires and received a thorough debriefing.

### Imaging details

#### Image acquisition

Participants were tested using a Philips Achieva 3T MRI scanner and a 32-channel SENSE headcoil. A survey scan was made for spatial planning of the subsequent scans. Following the survey scan, a 3-min structural T1-weighted scan was acquired using 3D fast field echo (TR: 82 ms, TE: 38 ms, flip angle: 8°, FOV: 240 × 188 mm, in-plane resolution 240 × 188, 220 slices acquired using single-shot ascending slice order and a voxel size of 1.0 × 1.0 × 1.0 mm). After the T1-weighted scan, functional T2*-weighted sequences were acquired using single shot gradient echo, echo planar imaging (TR = 2000 ms, TE = 27.63 ms, flip angle: 76.1°, FOV: 240 × 240 mm, in-plane resolution 64 × 64, 37 slices with ascending acquisition, slice thickness 3 mm, slice gap 0.3 mm, voxel size 3 × 3 × 3 mm), covering the entire brain. For the functional run, 495 volumes were acquired. After the functional run, a “B0” fieldmap scan (based on the phase difference between two consecutive echos) was acquired using 3D fast field echo (TR: 11 ms, TE: 3ms and 8ms, flip angle: 8°, FOV: 256 × 208, in-plane resolution 128 × 104, 128 slices).

#### Preprocessing

Results included in this manuscript come from preprocessing performed using FMRIPREP version 1.0.0 (Esteban et al., 2019; Esteban et al., 2017), a Nipype (Gorgolewski et al., 2011; Gorgolewski et al., 2017) based tool. Each T1 weighted volume was corrected for bias field using N4BiasFieldCorrection v2.1.0 (Tustison et al., 2010) and skullstripped using antsBrainExtraction.sh v2.1.0 (using OASIS template). Cortical surface was estimated using FreeSurfer v6.0.0 (Dale, Fischl, & Sereno, 1999). The skullstripped T1w volume was segmented (using FSL FAST; Zhang, Brady, & Smith, 2001) and coregistered to the skullstripped ICBM 152 Nonlinear Asymmetrical template version 2009c (Fonov et al., 2009) using nonlinear transformation implemented in ANTs v2.1.0 (Avants, Epstein, Grossman, & Gee., 2008).

Functional data was motion corrected using MCFLIRT v5.0.9 (Jenkinson, Bannister, Brady, & Smith, 2002). Distortion correction was performed using phase-difference fieldmaps processed with FUGUE (Jenkinson, 2003; FSL v5.0.9). This was followed by co-registration to the corresponding T1w using boundary-based registration (Greve & Fischl, 2009) with 9 degrees of freedom, using bbregister (FreeSurfer v6.0.0). Motion correcting transformations, field distortion correcting warp, BOLD-to-T1w transformation and T1w-to-template (MNI) warp were concatenated and applied in a single step using antsApplyTransforms (ANTs v2.1.0) using Lanczos interpolation.

Many internal operations of FMRIPREP use Nilearn (Abraham et al., 2014), principally within the BOLD-processing workflow. For more details of the pipeline see http://fmriprep.readthedocs.io/en/1.0.0/workflows.html.

#### First-level analysis

We modeled the participants’ preprocessed time series in a “first-level” GLM using *FSL FEAT* (Woolrich, Ripley, Brady, & Smith, 2001; FSL v6.0.0). The first-level modeling procedure was exactly the same for the participants in the active choice and passive viewing condition. As predictors, we included regressors for both the induction phase (i.e., the written description) and the relief phase (i.e., the full image). We separated trials with positive descriptions/images from trials with negative descriptions/images and separated trials in which participants saw the full version of the image from trials in which they saw a blurred version of the image. Note that in the active choice condition participants chose to see the full or blurred image, whereas in the passive viewing condition it was predetermined whether participants saw the full or blurred image. The final model held eight predictors: 2 (phase: induction vs. relief) × 2 (seen: full image vs. blurred image) × 2 (valence: negative vs. positive). If participants did not have any blurred image trials, the associated predictors were left out. Additionally, we added a single predictor for the actual decision (i.e., modelled at the onset the button press) and six motion predictors based on the estimated motion correction parameters.

Before model estimation, we applied a high-pass filter (σ = 50 seconds) and spatially smoothed the data (FWHM = 5 mm.). Standard prewhitening, as implemented in *FSL*, was applied. First-level contrasts only involved predictors associated with full image trials; that is, predictors associated with blurred image trials were not used for further analysis. For the remaining four predictors of interest — 2 (phase | full image) × 2 (valence | full image) — we defined contrasts against baseline, i.e., *β*_predictor_ ≠ 0, and valence contrasts, i.e., (*β*_neg | induction_ − *β*_pos | induction_) ≠ 0 and (*β*_neg | relief_ − *β*_pos | relief_) ≠ 0.

#### ROI-based group analysis

We tested two confirmatory hypotheses in this ROI-based group analysis, separately for the induction and relief phase:

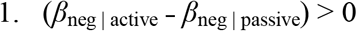

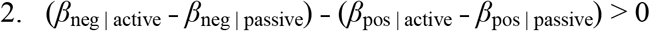

Note that the parameters (e.g., *β*_neg | active_) reflect the average of the first-level parameters (e.g., *β*_neg_) for a particular condition (e.g., active choice). As such, we tested four different group-level contrasts — 2 (phase) × 2 (hypothesis) — across two ROIs (striatum and IFG) in our group-level model.

For these confirmatory ROI-based group analyses, we used nonparametric permutation-based inference in combination with Threshold-Free Cluster Enhancement (TFCE; Smith & Nichols, 2009) as implemented in *FSL randomise* (Winkler, Ridgway, Webster, Smith, & Nichols, 2014). We ran *randomise* with 5000 permutations, corrected for multiple comparisons using the *maximum statistic* method (the method’s default multiple comparison correction procedure), and thresholded voxelwise results at *p* < 0.025 (correction for two ROIs). Note that this analysis allows for voxel-wise inference (i.e., no cluster-based correction is used).

In these ROI-based analyses, we restricted the analysis to voxels within two a-priori specified ROIs: bilateral striatum and bilateral inferior frontal gyrus (IFG). The ROIs are based on the Harvard-Oxford Subcortical Atlas (striatum; caudate, putamen and nucleus accumbens) and the Harvard-Oxford Cortical Atlas (IFG; pars opercularis and pars triangularis) with a threshold for probabilistic ROIs > 0 (Craddock et al., 2012).

#### Whole-brain group analysis

In addition to the confirmatory ROI-based analysis, we conducted an exploratory whole-brain group-analysis. Besides the two hypotheses mentioned in the previous section, we tested the following hypotheses, again for both the induction and relief phase:

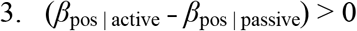

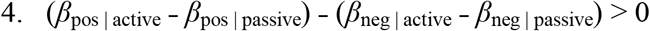

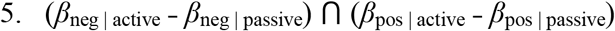

The ⋂ symbol in hypothesis 5 represents a conjunction analysis between two contrasts. For these exploratory whole-brain group analyses, we used *FSL FEAT* (Woolrich, Behrens, Beckmann, Jenkinson, & Smith, 2004) with a *FLAME1* mixed-effects model and automatic outlier detection (Woolrich, 2008). Resulting brain maps were thresholded with cluster-based correction (Worsley, 2001) using an initial (one-tailed) voxel-wise *p*-value cutoff of 0.005 (corresponding to a *z*-value above 2.576) and a cluster-wise significance level of 0.05. For the conjunction analysis (hypothesis 5), we used the *minimum statistic* approach (Nichols, Brett, Andersson, Wager, & Poline, 2005) in combination with cluster-based correction using the same cutoff and significance value as for the other two (non-conjunction based) hypotheses.

#### Further exploratory analyses

To aid interpretation of the results, we “decoded” the brain maps resulting from the whole-brain analysis using *Neurosynth* (Yarkoni et al., 2011; analyzed on March 4, 2019). In Supplementary Table 1, we list the ten Neurosynth terms (excluding anatomical terms) with the highest overall spatial correlation with our unthresholded brain maps (which are available on Neurovault, see below).

#### Code and data availability

All code used to preprocess, analyze, and plot the data is available from the project’s Github repository: https://github.com/lukassnoek/MorbidCuriosityFMRI. Unthresholded whole-brain group-level statistics maps are available from Neurovault (Gorgolewski et al., 2015): https://identifiers.org/neurovault.collection:5591. Much of this study’s code involves functionality from the *nilearn* Python package for neuroimaging analysis and visualization (Abraham et al., 2014).

## Supplementary materials

### Supplementary table S1

**Table S1.**
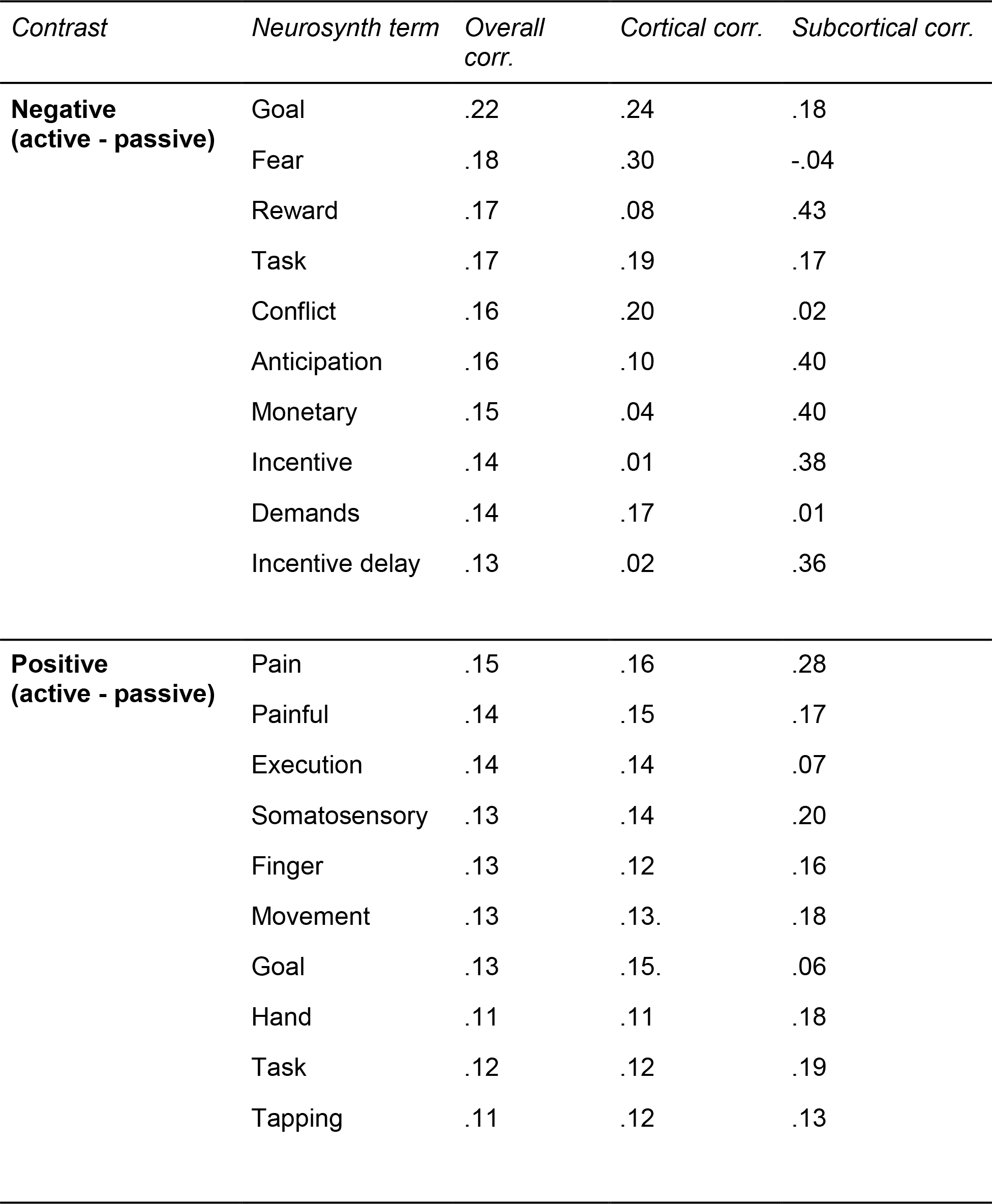
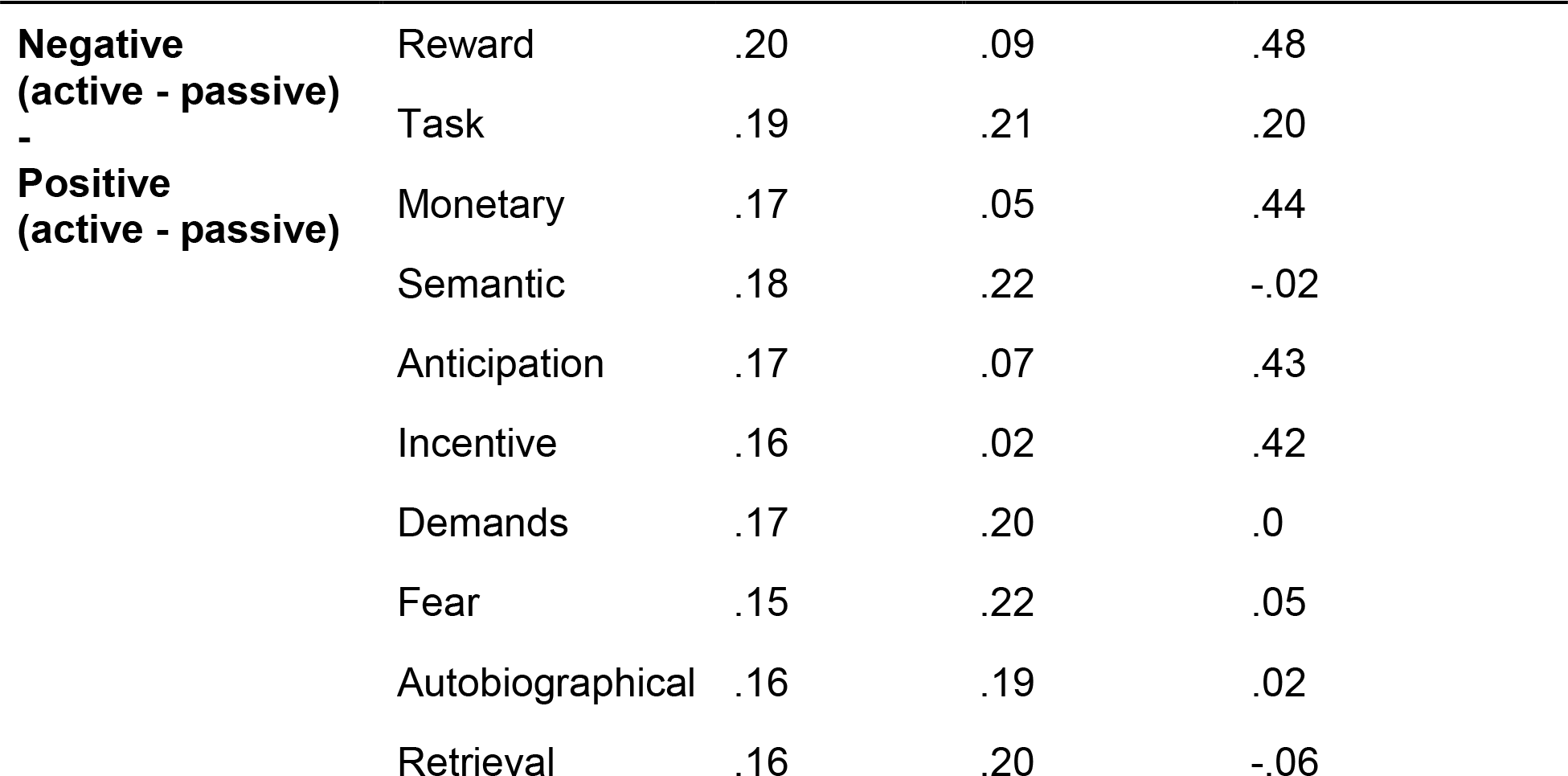
List of Neurosynth terms associated with the results of the exploratory whole-brain analyses for multiple contrasts. The ten terms with the highest spatial correlation with the whole-brain maps (excluding anatomical terms) are reported.

### Supplementary table S2

**Table S2.** Cluster statistics and associated brain regions from the exploratory whole-brain analysis of the contrast negative _active - passive_. The *X*, *Y*, and *Z* coordinates refer to MNI152 (2mm) space. The regions are taken from the Harvard-Oxford (sub)cortical atlas (Craddock et al., 2012) and voxels are assigned to regions based on their maximum probability across all ROIs within the atlas. *K* refers to the number of voxels within a particular region.

| Cluster nr. | Cluster size | Cluster max. | X | Y | Z | Region | K | Max. |
| --- | --- | --- | --- | --- | --- | --- | --- | --- |
| 1 | 12024 | 6.41 | 34 | 28 | -6 | Left Frontal Pole | 997 | 4.99 |
|  |  |  |  |  |  | Right Frontal Orbital Cortex | 870 | 6.41 |
|  |  |  |  |  |  | Left Frontal Orbital Cortex | 802 | 5.49 |
|  |  |  |  |  |  | Right Frontal Pole | 743 | 5.19 |
|  |  |  |  |  |  | Right Inferior Frontal Gyrus, pars triang. | 647 | 5.33 |
|  |  |  |  |  |  | Left Temporal Pole | 640 | 4.64 |
|  |  |  |  |  |  | Left Insular Cortex | 634 | 6.23 |
|  |  |  |  |  |  | Right Temporal Pole | 616 | 4.75 |
|  |  |  |  |  |  | Right Insular Cortex | 381 | 5.94 |
|  |  |  |  |  |  | Left Inferior Frontal Gyrus, pars operc. | 373 | 4.53 |
|  |  |  |  |  |  | Left Frontal Operculum Cortex | 365 | 5.37 |
|  |  |  |  |  |  | Left Inferior Frontal Gyrus, pars triang. | 324 | 5.49 |
|  |  |  |  |  |  | Right Frontal Operculum Cortex | 287 | 5.21 |
|  |  |  |  |  |  | Right Inferior Frontal Gyrus, pars opercularis | 218 | 4.48 |
|  |  |  |  |  |  | Left Middle Frontal Gyrus | 192 | 3.80 |
|  |  |  |  |  |  | Right Middle Frontal Gyrus | 102 | 3.50 |
|  |  |  |  |  |  | Right Middle Temporal Gyrus, anterior | 39 | 3.63 |
|  |  |  |  |  |  | Left Subcallosal Cortex | 39 | 4.23 |
|  |  |  |  |  |  | Left Precentral Gyrus | 27 | 3.22 |
|  |  |  |  |  |  | Left Central Opercular Cortex | 27 | 3.12 |
|  |  |  |  |  |  | Right Thalamus | 462 | 4.76 |
|  |  |  |  |  |  | Brain-Stem, left part | 447 | 4.19 |
|  |  |  |  |  |  | Left Thalamus | 446 | 4.40 |
|  |  |  |  |  |  | Left Caudate | 257 | 4.39 |
|  |  |  |  |  |  | Right Caudate | 239 | 4.95 |
|  |  |  |  |  |  | Right Putamen | 211 | 4.53 |
|  |  |  |  |  |  | Left Putamen | 173 | 4.88 |
|  |  |  |  |  |  | Brain-Stem, right part | 153 | 4.34 |
|  |  |  |  |  |  | Left Accumbens | 61 | 4.16 |
|  |  |  |  |  |  | Right Pallidum | 34 | 3.63 |
|  |  |  |  |  |  | Right Accumbens | 29 | 4.31 |
|  |  |  |  |  |  | Right Amygdala | 26 | 3.69 |
| 2 | 7133 | 6.17 | 8 | 30 | 26 | Right Paracingulate Gyrus | 974 | 5.95 |
|  |  |  |  |  |  | Right Cingulate Gyrus, anterior division | 897 | 6.17 |
|  |  |  |  |  |  | Left Paracingulate Gyrus | 835 | 5.99 |
|  |  |  |  |  |  | Left Superior Frontal Gyrus | 832 | 4.73 |
|  |  |  |  |  |  | Left Cingulate Gyrus, anterior division | 797 | 5.89 |
|  |  |  |  |  |  | Right Superior Frontal Gyrus | 646 | 4.67 |
|  |  |  |  |  |  | Right Frontal Pole | 443 | 4.25 |
|  |  |  |  |  |  | Left Juxtapositional Lobule Cortex | 360 | 5.04 |
|  |  |  |  |  |  | Left Postcentral Gyrus | 287 | 4.15 |
|  |  |  |  |  |  | Left Precentral Gyrus | 283 | 4.08 |
|  |  |  |  |  |  | Right Juxtapositional Lobule Cortex | 281 | 4.78 |
|  |  |  |  |  |  | Left Frontal Pole | 179 | 4.11 |
|  |  |  |  |  |  | Left Middle Frontal Gyrus | 124 | 4.03 |
|  |  |  |  |  |  | Left Superior Parietal Lobule | 42 | 3.64 |
|  |  |  |  |  |  | Right Cingulate Gyrus, posterior division | 40 | 3.41 |
|  |  |  |  |  |  | Right Frontal Medial Cortex | 23 | 3.13 |
| 3 | 534 | 5.05 | -34 | -46 | -24 | Left Temporal Fusiform Cortex, posterior | 95 | 3.98 |
|  |  |  |  |  |  | Left Temporal Occipital Fusiform Cortex | 76 | 5.05 |
| 4 | 401 | 4.13 | 20 | -50 | 0 | Right Lingual Gyrus | 182 | 4.13 |
|  |  |  |  |  |  | Right Intracalcarine Cortex | 119 | 3.77 |
|  |  |  |  |  |  | Right Supracalcarine Cortex | 38 | 3.47 |
|  |  |  |  |  |  | Right Cingulate Gyrus, posterior division | 35 | 3.78 |
| 5 | 375 | 4.70 | -16 | -50 | -2 | Left Lingual Gyrus | 115 | 4.70 |
|  |  |  |  |  |  | Left Cingulate Gyrus, posterior division | 111 | 4.05 |
|  |  |  |  |  |  | Left Precuneous Cortex | 61 | 3.97 |

### Supplementary table S3

**Table S3.**
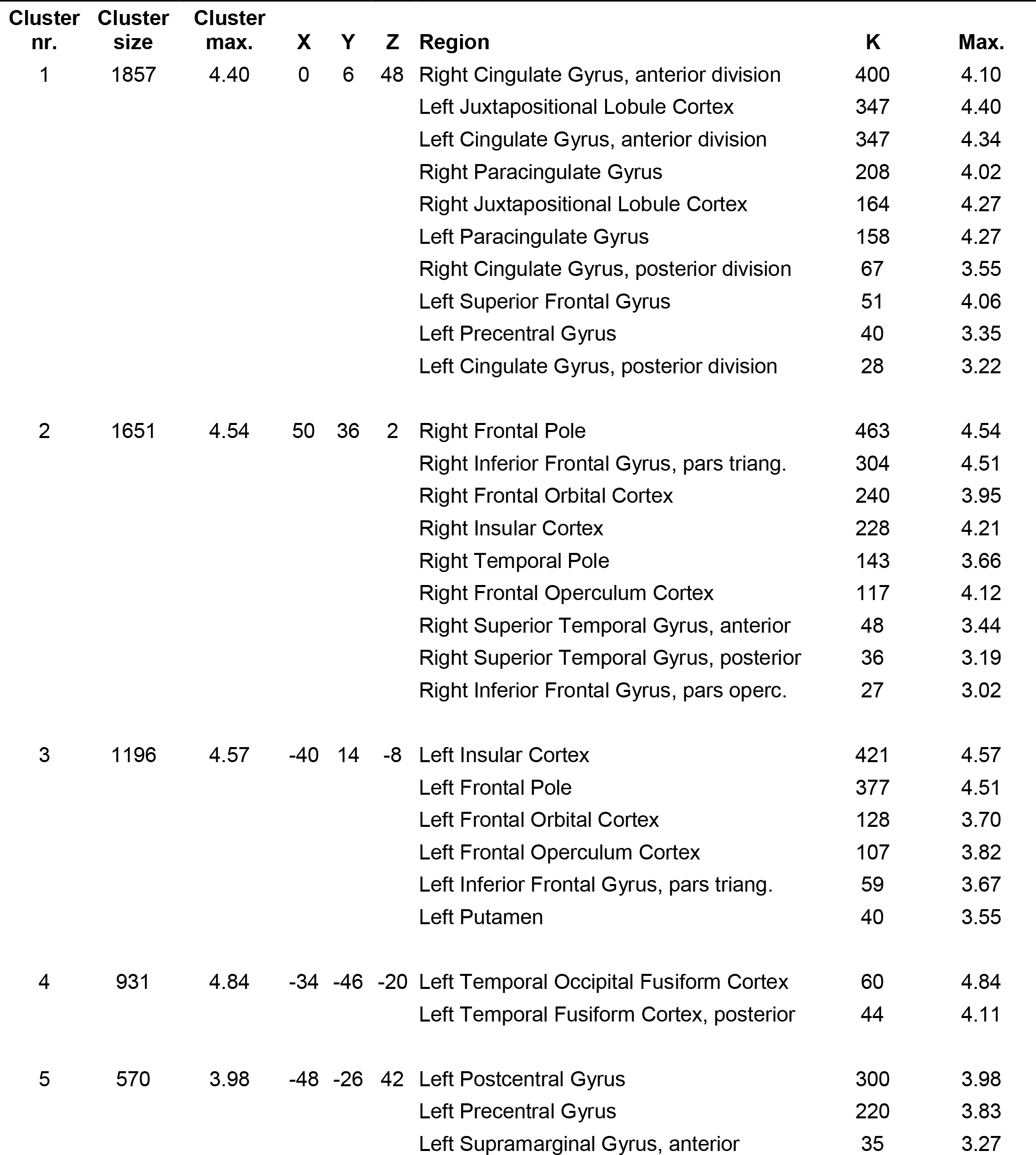
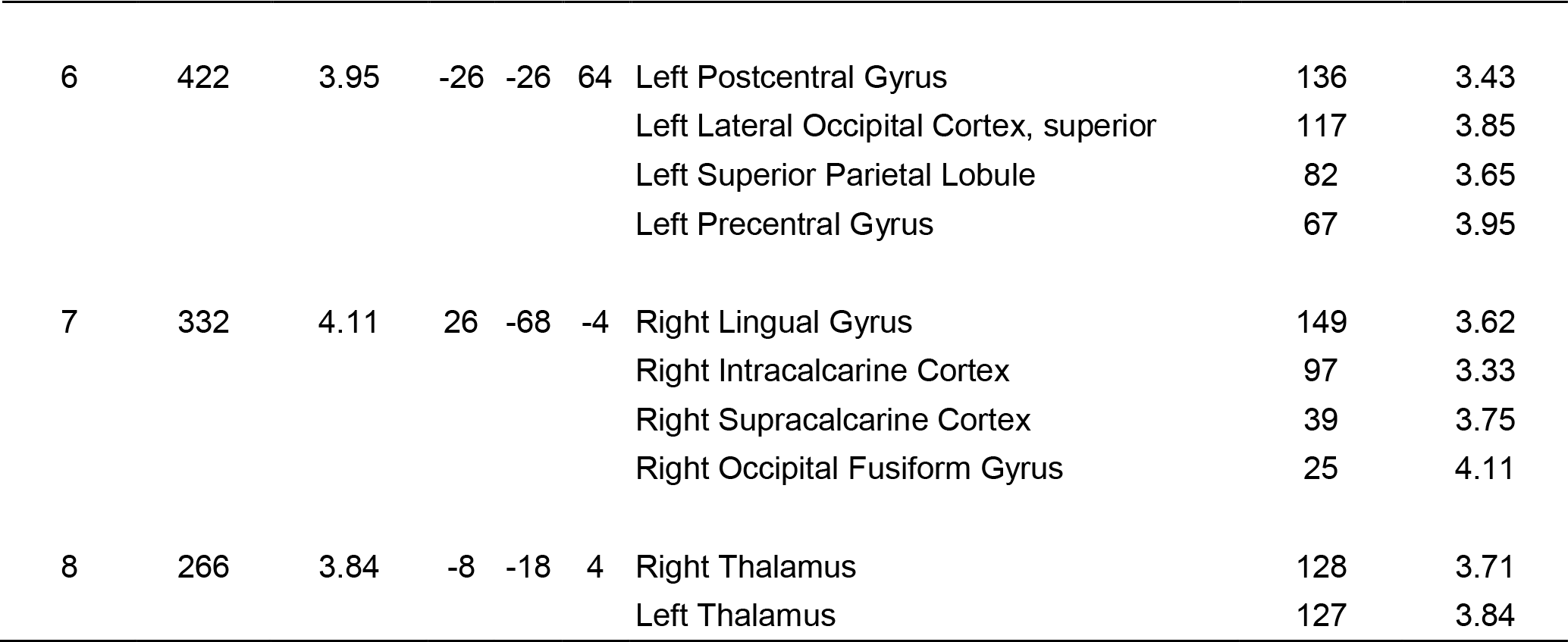
Cluster statistics and associated brain regions from the exploratory whole-brain analysis of the contrast positive _active - passive_. The *X*, *Y*, and *Z* coordinates refer to MNI152 (2mm) space. The regions are taken from the Harvard-Oxford (sub)cortical atlas (Craddock et al., 2012) and voxels are assigned to regions based on their maximum probability across all ROIs within the atlas. *K* refers to the number of voxels within a particular region.

| Cluster nr. | Cluster size | Cluster max. | X | Y | Z | Region | K | Max. |
| --- | --- | --- | --- | --- | --- | --- | --- | --- |
| 1 | 1857 | 4.40 | 0 | 6 | 48 | Right Cingulate Gyrus, anterior division | 400 | 4.10 |
|  |  |  |  |  |  | Left Juxtapositional Lobule Cortex | 347 | 4.40 |
|  |  |  |  |  |  | Left Cingulate Gyrus, anterior division | 347 | 4.34 |
|  |  |  |  |  |  | Right Paracingulate Gyrus | 208 | 4.02 |
|  |  |  |  |  |  | Right Juxtapositional Lobule Cortex | 164 | 4.27 |
|  |  |  |  |  |  | Left Paracingulate Gyrus | 158 | 4.27 |
|  |  |  |  |  |  | Right Cingulate Gyrus, posterior division | 67 | 3.55 |
|  |  |  |  |  |  | Left Superior Frontal Gyrus | 51 | 4.06 |
|  |  |  |  |  |  | Left Precentral Gyrus | 40 | 3.35 |
|  |  |  |  |  |  | Left Cingulate Gyrus, posterior division | 28 | 3.22 |
| 2 | 1651 | 4.54 | 50 | 36 | 2 | Right Frontal Pole | 463 | 4.54 |
|  |  |  |  |  |  | Right Inferior Frontal Gyrus, pars triang. | 304 | 4.51 |
|  |  |  |  |  |  | Right Frontal Orbital Cortex | 240 | 3.95 |
|  |  |  |  |  |  | Right Insular Cortex | 228 | 4.21 |
|  |  |  |  |  |  | Right Temporal Pole | 143 | 3.66 |
|  |  |  |  |  |  | Right Frontal Operculum Cortex | 117 | 4.12 |
|  |  |  |  |  |  | Right Superior Temporal Gyrus, anterior | 48 | 3.44 |
|  |  |  |  |  |  | Right Superior Temporal Gyrus, posterior | 36 | 3.19 |
|  |  |  |  |  |  | Right Inferior Frontal Gyrus, pars operc. | 27 | 3.02 |
| 3 | 1196 | 4.57 | -40 | 14 | -8 | Left Insular Cortex | 421 | 4.57 |
|  |  |  |  |  |  | Left Frontal Pole | 377 | 4.51 |
|  |  |  |  |  |  | Left Frontal Orbital Cortex | 128 | 3.70 |
|  |  |  |  |  |  | Left Frontal Operculum Cortex | 107 | 3.82 |
|  |  |  |  |  |  | Left Inferior Frontal Gyrus, pars triang. | 59 | 3.67 |
|  |  |  |  |  |  | Left Putamen | 40 | 3.55 |
| 4 | 931 | 4.84 | -34 | -46 | -20 | Left Temporal Occipital Fusiform Cortex | 60 | 4.84 |
|  |  |  |  |  |  | Left Temporal Fusiform Cortex, posterior | 44 | 4.11 |
| 5 | 570 | 3.98 | -48 | -26 | 42 | Left Postcentral Gyrus | 300 | 3.98 |
|  |  |  |  |  |  | Left Precentral Gyrus | 220 | 3.83 |
|  |  |  |  |  |  | Left Supramarginal Gyrus, anterior | 35 | 3.27 |
| 6 | 422 | 3.95 | -26 | -26 | 64 | Left Postcentral Gyrus | 136 | 3.43 |
|  |  |  |  |  |  | Left Lateral Occipital Cortex, superior | 117 | 3.85 |
|  |  |  |  |  |  | Left Superior Parietal Lobule | 82 | 3.65 |
|  |  |  |  |  |  | Left Precentral Gyrus | 67 | 3.95 |
| 7 | 332 | 4.11 | 26 | -68 | -4 | Right Lingual Gyrus | 149 | 3.62 |
|  |  |  |  |  |  | Right Intracalcarine Cortex | 97 | 3.33 |
|  |  |  |  |  |  | Right Supracalcarine Cortex | 39 | 3.75 |
|  |  |  |  |  |  | Right Occipital Fusiform Gyrus | 25 | 4.11 |
| 8 | 266 | 3.84 | -8 | -18 | 4 | Right Thalamus | 128 | 3.71 |
|  |  |  |  |  |  | Left Thalamus | 127 | 3.84 |

### Supplementary Table S4

**Table S4.**
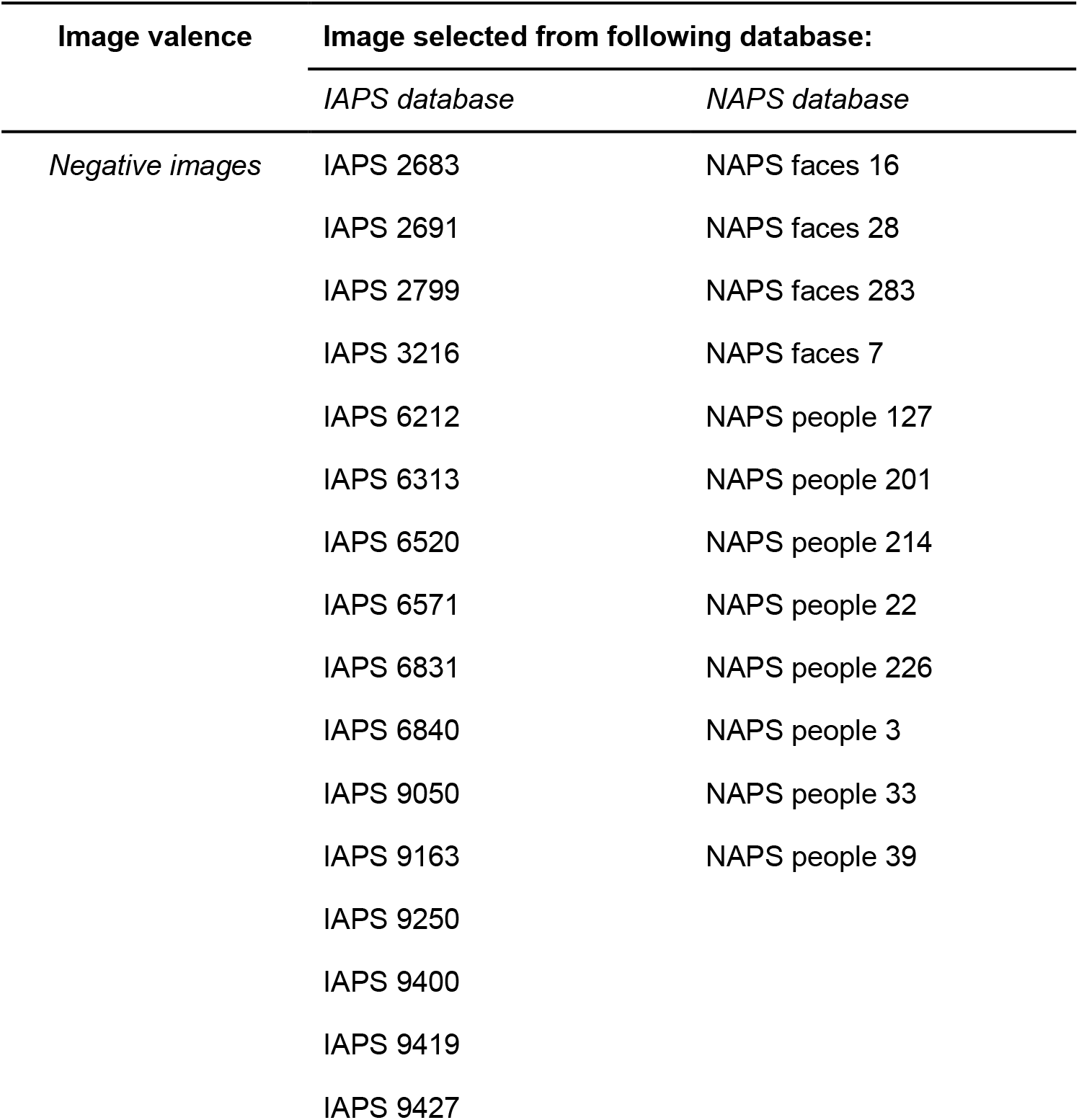
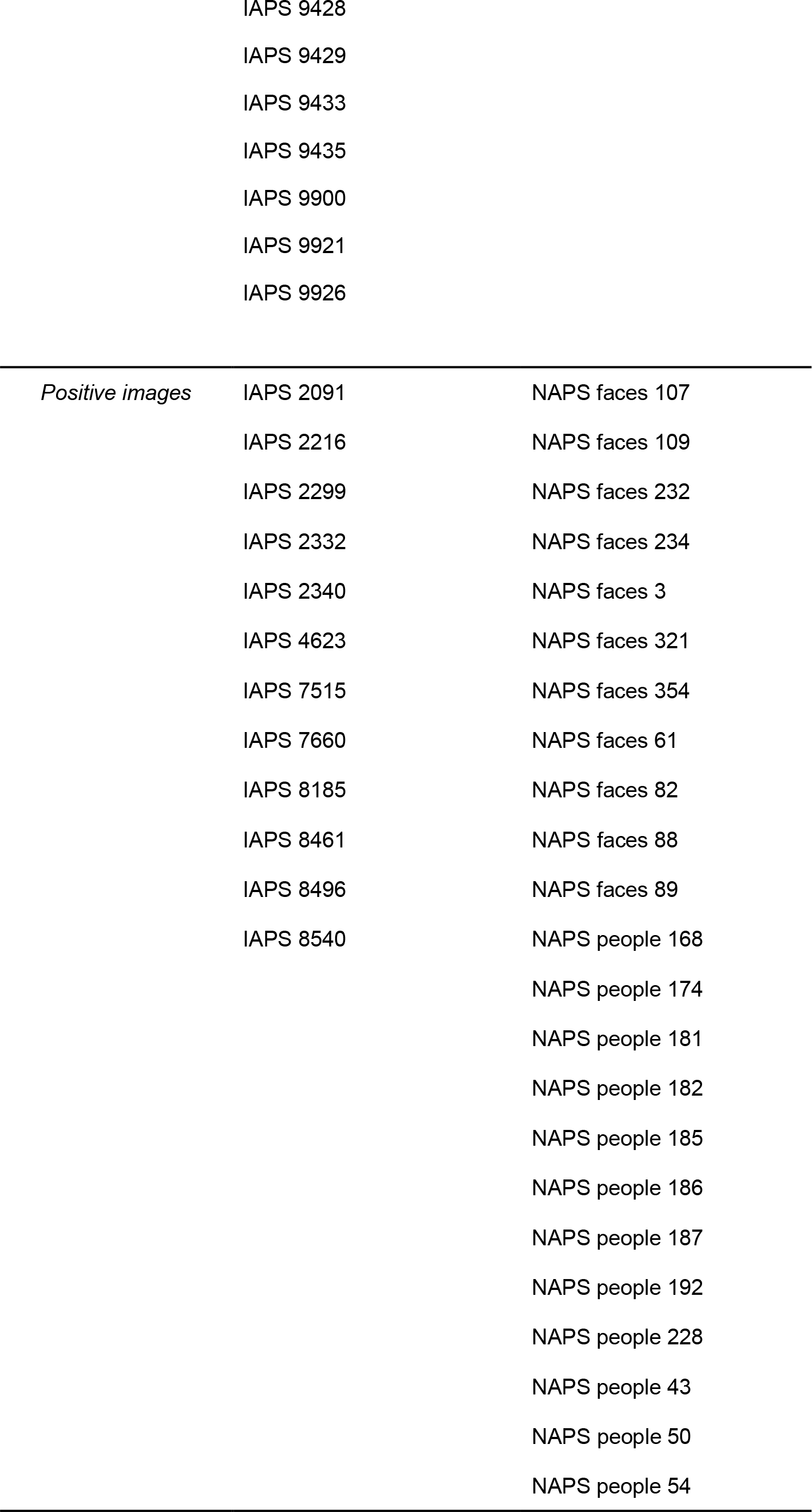
Stimulus codes for the images used in the choice task. Images were taken from both the IAPS (Lang, Bradley, & Cuthbert, 1997) and NAPS (Marchewka, Żurawski, Jednoróg, & Grabowska, 2014) database.

### Supplementary Figure S1

**Figure S1.**
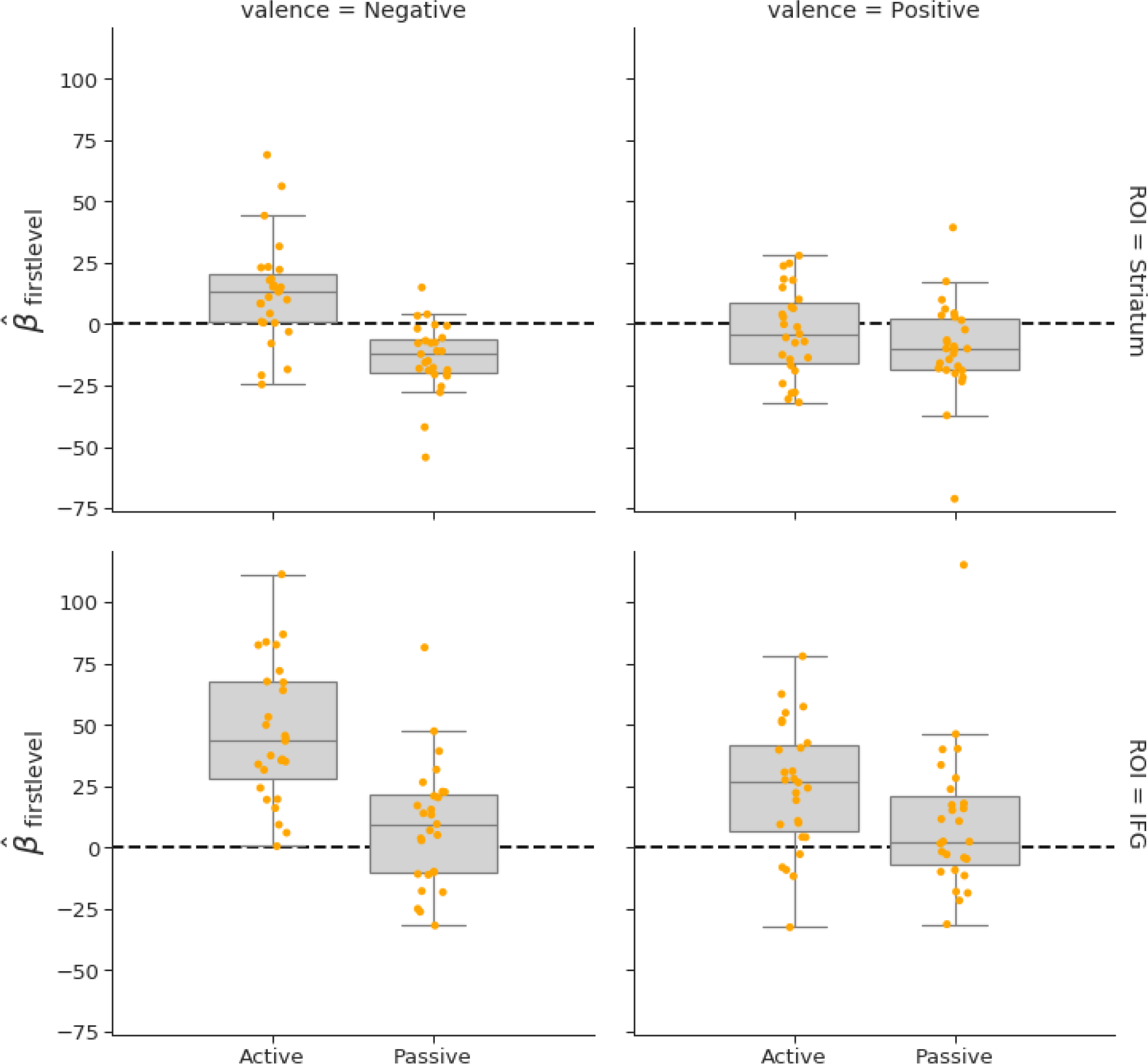
Subplots of individual regressors from the significant voxels in the confirmatory contrast negative _active - passive_ - positive _active - passive_ in the induction-phase. These plots show the direction of the effects. Plots are averaged over all significant voxels within each ROI (striatum in upper plots, IFG in lower plots), separately for the negative trials (left plots) and positive trials (right plots) with subplots for the active choice and passive viewing condition. Dots represent the participant-specific ROI-average parameter estimate from the first-level analysis. The horizontal line in the boxplots represents the median and the whiskers represent the interquartile range. Note that this figure is only meant to show the directionality of the effects, not their statistical significance (as the ROIs itself only contain voxels that were significant in the group-analysis).

